# Impact of acoustic property uncertainty on transcranial ultrasound simulation

**DOI:** 10.64898/2026.10.30.763065

**Authors:** Julian Q. Kosciessa, Taylor Webb, Eleanor Martin, Brad Treeby, Kim Butts Pauly, Lennart Verhagen, Samuel Pichardo

## Abstract

Subject-specific simulations of wave propagation are commonly used to plan and report acoustic and thermal exposure during transcranial ultrasound (TUS) interventions. Their accuracy depends on acoustic and thermal tissue property assumptions, including sound speed, density, and attenuation. Reported property measurements vary widely across the literature, precluding consensus point estimates for simulation. How this property uncertainty affects estimated exposure has not been quantified. We compile literature-based bounds for seven acoustic and thermal properties of skull and brain tissue and assess their individual impact on simulated intensity and temperature rise using two independent simulation platforms (PRESTUS/k-Wave and BabelBrain). Variations in skull attenuation and absorption produced the largest outcome uncertainty: peak intensity in the brain varied roughly threefold across the reported range of skull attenuation alone, while absorption most strongly affected temperature rise within the skull. The direction of each parameter’s effect was consistent across platforms and, with one exception, across skull thickness. This allowed bounds to be combined into setups that bracket transmission and temperature rise. Compared to these extremes, default property sets of three simulation toolboxes (k-Plan, BabelBrain, PRESTUS) yielded similar brain intensity but divergent skull heating, driven mainly by differing assumptions on skull absorption and heat capacity. These results indicate that uncertainty in skull acoustic properties is a major source of variability in TUS exposure estimates, and that refined measurements of these properties, together with explicit reporting of exposure uncertainty, are needed to support safe, effective, and reproducible transcranial ultrasound planning.

## 1. Introduction

Transcranial ultrasound stimulation (TUS) is a key technology used to non-invasively target neural activity in humans, with growing interest in neuromodulation and therapeutic applications (Murphy et al., 2025). Ensuring safety and efficacy of TUS applications requires reliable estimation of the acoustic and thermal exposure delivered through head tissues. Because direct in vivo measurement of intracranial acoustic pressure and temperature rise is not feasible, numerical simulation of ultrasound propagation across heterogeneous head tissues has become the prevalent approach for estimating exposure (Angla et al., 2023) and determining risk prior to stimulation (Aubry et al., 2025; Martin et al., 2024; Murphy et al., 2025).

Multiple platforms now simulate the intracranial ultrasound field and temperature rise from individual head geometry and composition. In practice, however, their outputs differ: a recent comparison of two widely used simulation platforms (k-Plan and BabelBrain) reported variable in-situ intensity and temperature rise estimates for the same individuals, trajectories, and sonication protocols (Brandts et al., 2026). Different solvers have been shown to converge to the same results when given identical inputs and tissue properties (Aubry et al., 2022), suggesting that numerical methods are not the primary source of variation. Instead, modeling assumptions such as the acoustic and thermal properties assigned to head tissues introduce uncertainty in estimates of TUS exposure (Robertson, Martin, et al., 2017).

Sound speed, density, and attenuation determine ultrasound propagation through head tissues, and thereby the magnitude and spatial distribution of pressure and intensity at the target. Specific heat capacity, thermal conductivity, and perfusion additionally determine the temperature rise predicted by bioheat models (e.g., the Pennes equation; Pennes, 1948). Beyond expected intra- and interindividual variability, reported property values reflect a heterogeneous literature, with differences in specimen type, age, bone composition (Fry & Barger, 1978; Goss et al., 1979), experimental setup, sample preparation (e.g., ex vivo vs. in vivo), and ultrasound frequency.

The impact of tissue property uncertainty needs systematic quantification to ensure consistency across TUS simulation platforms. This study, led by the Acoustic Properties working group of the International Transcranial Ultrasonic Stimulation Safety and Standards Consortium (ITRUSST), addresses this gap. We compile the empirically reported range of skull and brain acoustic and thermal properties from the literature, use a sensitivity analysis in skull and brain benchmark models to identify which parameters contribute most to exposure uncertainty, and combine parameter bounds to estimate composite exposure uncertainty.

## 2. Methods

### 2.1 Literature review of skull and brain acoustic parameters

Tissue parameter values were derived from a targeted collection of the primary measurement literature for skull and brain acoustic (sound speed, density, attenuation) and thermal properties (thermal conductivity, specific heat capacity, perfusion, and absorption). This collection is shown in Supp. Table S1 and forms the foundation of an evolving ITRUSST Tissue Properties Database (Kosciessa, Webb, et al., 2026), available at https://itrusst.github.io/itrusst_properties/.

For inclusion in the collection, we targeted values that describe intrinsic properties of adult, healthy human skull and brain tissues that could be traced to a primary measurement. For skull acoustic properties, we included measurements within 0.2 to 1.03 MHz, the frequency range typically used in TUS. However, brain attenuation has been measured almost exclusively above 1 MHz, so these values were retained and extrapolated (see below). Because human cranial data are sparse, particularly for thermal properties, some values from non-human and non-cranial samples were retained. Where a study reported several specimens of the same tissue and preparation, we listed the study’s mean where available; specimens that differed in preparation, site or layer, and studies without a reported mean, were listed individually. We excluded values that (i) had no traceable source or specimen or were stated as arbitrary; (ii) misreported their primary source in a secondary compilation; or (iii) reported transmission or insertion loss, or included fitted reflection losses, instead of attenuation. Supp. Table S1 lists exclusion reasons. Attenuation coefficients were scaled to 0.5 MHz using the reported or study-fitted frequency exponent b, with b = 1 assumed where only a single frequency was available. Retained minima and maxima defined LOW and HIGH bounds (Table 1).

**Table 1.**
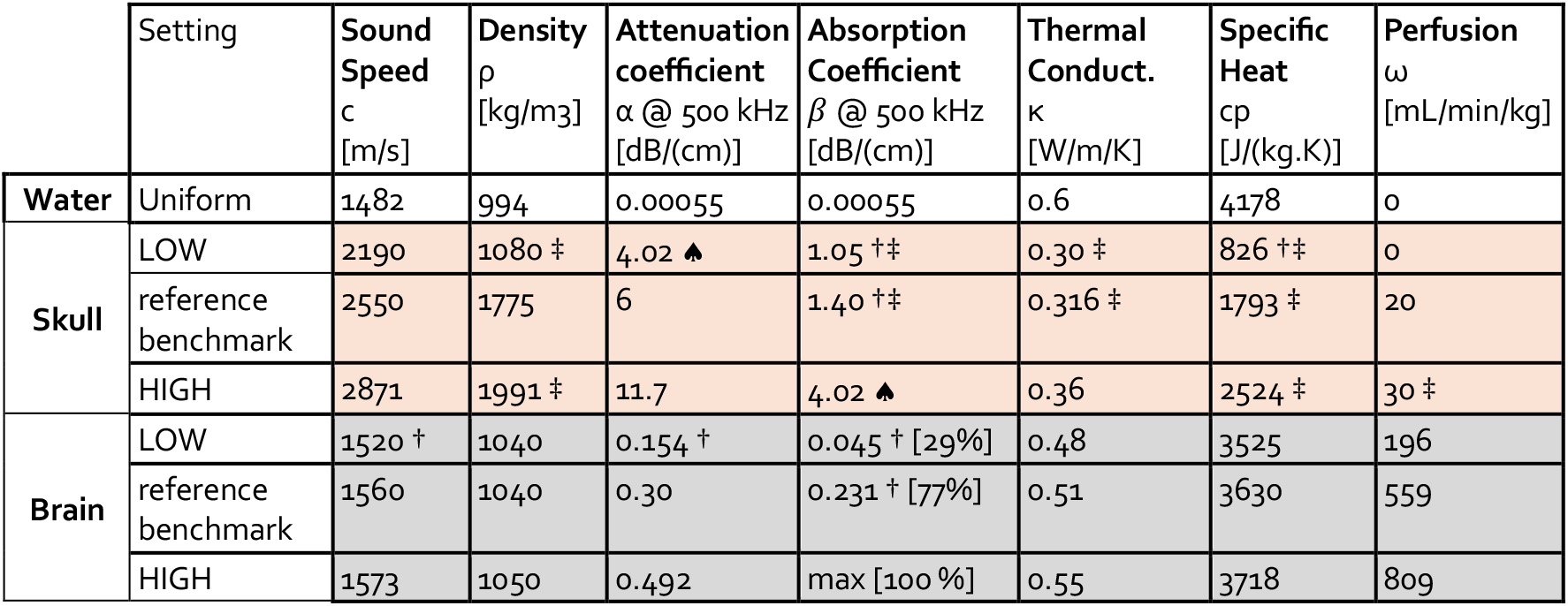
Acoustic and thermal property values in the sensitivity analysis. LOW and HIGH correspond to the minimum and maximum values compiled in Supp. Table S1 (v0.1.0 of the ITRUSST Tissue Properties Database). Reference benchmark values should not be interpreted as optimal point estimates (see methods). † Value from a non-human sample. ‡ Value from non-skull bone. ♠ Low skull attenuation used as a proxy for maximal skull absorption.

|  | Setting | **Sound Speed**<br>c<br>[m/s] | **Density**<br>ρ<br>[kg/m3] | **Attenuation coefficient**<br>α @ 500 kHz<br>[dB/(cm)] | **Absorption Coefficient**<br>*β* @ 500 kHz<br>[dB/(cm)] | **Thermal Conduct.**<br>κ<br>[W/m/K] | **Specific Heat**<br>cp<br>[J/(kg.K)] | **Perfusion**<br>ω<br>[mL/min/kg] |
|---|---|---|---|---|---|---|---|---|
| **Water** | Uniform | 1482 | 994 | 0.00055 | 0.00055 | 0.6 | 4178 | 0 |
| **Skull** | LOW | 2190 | 1080 ‡ | 4.02 ♠ | 1.05 †‡ | 0.30 ‡ | 826 †‡ | 0 |
|  | reference benchmark | 2550 | 1775 | 6 | 1.40 †‡ | 0.316 ‡ | 1793 ‡ | 20 |
|  | HIGH | 2871 | 1991 ‡ | 11.7 | 4.02 ♠ | 0.36 | 2524 ‡ | 30 ‡ |
| **Brain** | LOW | 1520 † | 1040 | 0.154 † | 0.045 † [29%] | 0.48 | 3525 | 196 |
|  | reference benchmark | 1560 | 1040 | 0.30 | 0.231 † [77%] | 0.51 | 3630 | 559 |
|  | HIGH | 1573 | 1050 | 0.492 | max [100 %] | 0.55 | 3718 | 809 |

Ultrasound loses energy in the skull through reflection at interfaces, absorption (conversion to heat), and scattering by the porous inner structure. The compiled attenuation coefficients are intended to capture absorption and scattering inside the bone, not interface reflections, although some values may include partial reflection loss. Thermal simulations have often modeled absorption as a fraction of attenuation, with large variability among toolboxes, up to full absorption in k-Plan (k-Plan v1.2.1; Brainbox Ltd., 2024). When coupled to high attenuation, such fractions can assume absorption far beyond reported measurements. We therefore chose to model absorption using the absorption coefficient as a semi-independent parameter. As only two skull absorption measurements were identified and our models do not capture shear-wave contributions to skull heating, we used the low estimate of skull attenuation as a conservative proxy for high absorption (P. J. White et al., 2006).

Brain absorption was modeled as a fraction of brain attenuation, because low absorption coefficients combined with high attenuation would imply implausibly small fractions. Measured fractions remain sparse and variable: in bovine brain at 20–22 °C, composite gray/white matter absorption corresponded to 77–100% of the attenuation measured in mixed tissue (Lyons & Parker, 1988). In contrast, cat brain absorption at 37 °C (Goss et al., 1979) corresponds to approximately 20–30% of brain attenuation measured at the same temperature in humans (Kremkau et al., 1981), although this comparison spans studies and species. To cover this range, we divided the absorption coefficient by the attenuation coefficient at the same level, yielding fractions of 29% for LOW (0.045/0.154 dB/cm) and 77% at the reference level (0.231/0.30 dB/cm). For HIGH, we assumed 100% or complete absorption.

Acoustic reference values (sound speed, density, attenuation coefficient) for skull and brain follow an intercomparison benchmark (Aubry et al., 2022), averaging cortical and trabecular bone estimates for skull. Thermal reference values follow the IT’IS Foundation Tissue Properties Database v.5.0 (Baumgartner et al., 2025). The reference skull absorption coefficient was set to 1.40 dB/cm at 500 kHz (D. N. White et al., 1978). The brain absorption coefficient was set to 0.231 dB/cm at 500 kHz, the highest value reported in the (non-human) brain absorption literature (Lyons & Parker, 1988). This value was chosen over mid-range values given the sparsity of absorption measurements and the low attenuation measures in the dataset.

### 2.2 Model specifications

Acoustic and thermal simulations were performed in two configurations: (1) water & soft tissue and (2) water, skull, & soft tissue (Fig. 1a), modeled on a 120 × 70 mm two-dimensional axisymmetric grid at 0.5 mm isotropic resolution. A single skull compartment was chosen due to the unreliability of compartment-specific parameter values and the more complicated dependence of multi-compartment models on layer thickness and ultrasound frequency. A single-element focused ultrasound transducer with 64 mm aperture, 64 mm focal length (Aubry et al., 2022) was modeled at 500 kHz. The source amplitude was chosen to generate a free-field spatial-peak pulse-average intensity (I_SPPA_) of 60 W/cm^2^. The stimulation protocol consisted of 20 ms pulses at 5 Hz (10% duty cycle) delivered over an 80 s pulse train, followed by an 80 s post-stimulation window to assess thermal propagation to near steady-state.

**Figure 1.**
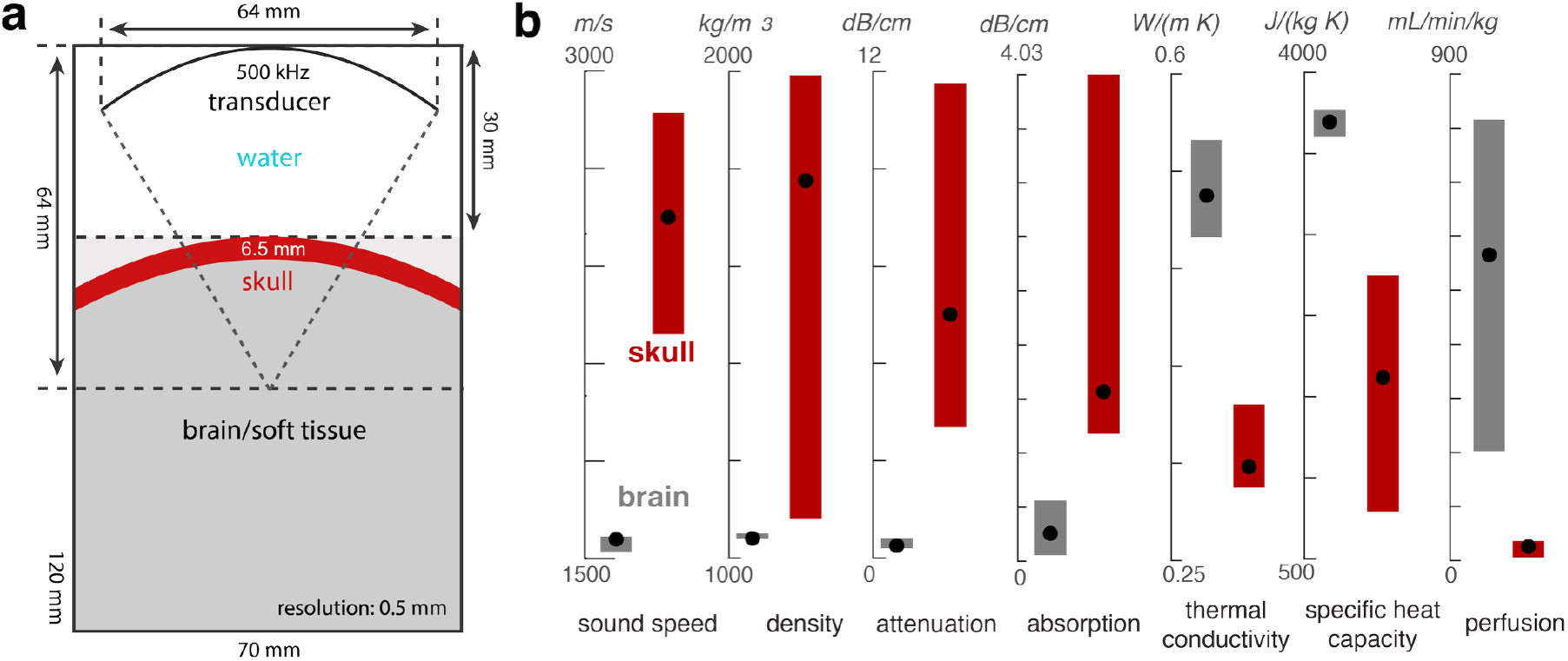
Benchmark setup. (a) Schematic of the two tissue configurations (model 1: brain; model 2: skull + brain). The light gray shading indicates the location of the soft tissue layer in model 1. (b) Acoustic and thermal property ranges of brain and skull tissues: black dots indicate reference values for each parameter (see Table 1).

In a sensitivity analysis, acoustic and thermal tissue parameters (sound speed, density, attenuation, absorption, specific heat capacity, thermal conductivity, and perfusion rate) were varied one-at-a-time between their LOW, HIGH, and reference levels, independently for brain and skull compartments. Remaining parameters were held constant at their reference levels. Estimated peak intensity in the brain (I_sppa_) and peak temperature rise (ΔT) in the skull and at the focus served as the primary outcome metrics.

Two independent simulation pipelines ran the benchmarks in parallel to assess cross-tool robustness of parameter impact. In PRESTUS (v0.6.1; Kosciessa, Chetverikov, et al., 2026), compressional wave propagation was simulated using k-Wave (v1.4.1; Treeby & Cox, 2010) on axisymmetric grids (kspaceFirstOrderAS; Treeby et al., 2020). An axisymmetric solver was used to increase computational efficiency with a 20 mm perfectly matched layer (PML). Tissue heating was modeled by numerically solving the Pennes bioheat equation (Pennes, 1948) via kWaveDiffusion (Treeby & Cox, 2010). The volumetric heat source was computed from the simulated peak pressure as Q = *β* p^2^/(ρc), where *β* is the tissue absorption coefficient at the source frequency, and was applied during pulses only. In BabelBrain, compressional wave propagation and viscoelastic skull interactions were simulated using a Rayleigh–Sommerfeld approach, viscoelastic wave propagation (BabelViscoFDTD), and bioheat-transfer-equation (BHTE) FDTD solver (v0.8.8; Pichardo, 2023). Code reproducing the setup and analyses is available at https://github.com/itrusst/itrusst_properties.

To test whether parameter effects on exposure generalize across skull thickness, we first varied skull thickness from 1 to 12 mm in 0.5 mm steps, with all tissue properties held at reference levels. The outer surface remained of fixed radius (75 mm, apex 30 mm from the transducer on axis) and the inner radius was reduced by the desired thickness. We applied sensitivity analysis for the skull thickness with maximal transmission for our setup to probe the robustness of parameter effects.

To investigate the joint uncertainty arising from parameter constellations, we identified the direction of independent parameter impact on acoustic intensity and skull heating and aligned minimum/maximum properties across all seven tissue properties to min/max I_sppa_ or skull heating. We compared these bounded setups to default tissue-parameter presets published for three toolboxes (PRESTUS v0.6.1, BabelBrain v0.8.8, and k-Plan v1.2.1) evaluated within the PRESTUS/k-Wave solver. These emulations do not capture other toolbox differences such as k-Plan’s CT-based density derivation in acoustic simulations.

## 3. Results

### 3.1 Substantial variability in literature-based parameter estimates

A targeted literature review compiled skull and brain property values from 38 sources (Fig. 1b; Supp. Table S1). Values from 6 sources were excluded partially or totally. The spread in values, which included measurements from human and non-human specimens, was wide for skull and brain alike: for example, at 0.5 MHz, skull attenuation varied approximately 3-fold, skull sound speed 1.3-fold, and skull density 1.8-fold. Measurements obtained were between 0.27 and 2.5 MHz for skull and mostly between 1 and 6 MHz for brain, so they had to be extrapolated to 0.5 MHz using frequency exponents that themselves varied between studies (approximately 0.8–1.5). Experimental methods differed considerably: skull measurements stem from one to seven ex vivo skulls per study, sampled at different anatomical sites and either as whole sections or as separated cortical and diploic layers. Specimen preparation ranged from fresh tissue to formalin-fixed and fully dried bone. Several compilation values, including all IT’IS skull attenuation values, derive from non-human models (bovine or porcine bone, often femur rather than skull). Thermal properties were informed by fewer measurements overall and came mostly from non-cranial or non-human samples. No measured values of human brain absorption were identified. These literature ranges defined the LOW and HIGH bounds (Table 1, Fig. 1b) for the benchmark simulations.

### 3.2 Robust individual parameter impact on computed in-situ intensity

We first varied each property one at a time between its LOW and HIGH bound and quantified the change in brain I_SPPA_ (Fig. 2). Brain property uncertainty had little effect on brain I_SPPA_, except for a minor effect of brain attenuation, whereas uncertainty about skull properties, chief among them skull attenuation, produced substantial transmission variation. Increasing skull attenuation reduced brain I_SPPA_, consistent with less energy reaching the focus. Across the literature range of skull attenuation alone, brain I_SPPA_ ranged from 4 to 11 W/cm^2^ for our protocol, a roughly threefold difference with all other parameters held constant. The direction of each parameter’s effect was reproduced in both pipelines. As the reference skull sound speed (2550 m/s) lies close to the transmission minimum for the 6.5 mm skull at 500 kHz, intensity at reference values fell below the LOW-to-HIGH range. Focal position and volume were largely robust to tissue property variations: sound speed and attenuation shifted the focus by at most 1 mm (Supp. Fig. 1).

**Figure 2.**
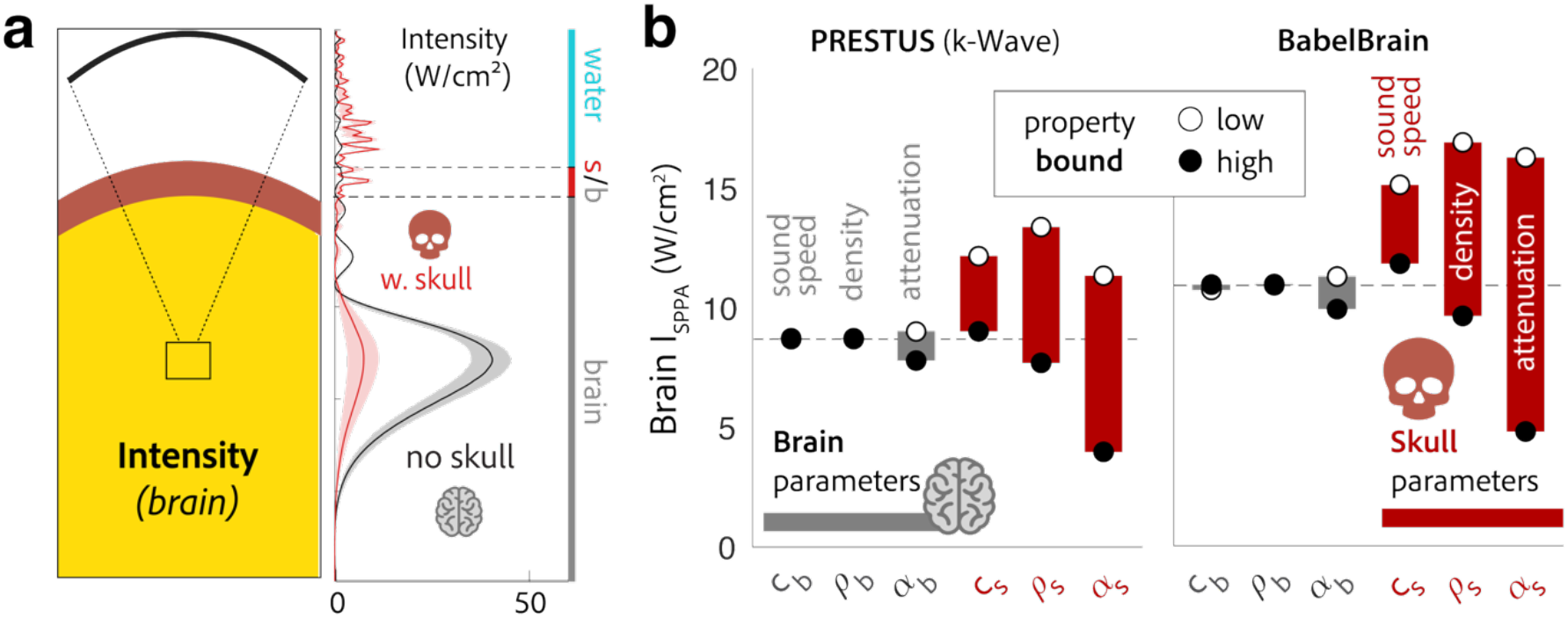
Impact of parameter uncertainty on computed in-situ focal intensity. (a) Benchmark schematic (left; yellow indicates the read-out region) and on-axis pulse-average intensity in PRESTUS (right) without (black) and with (red) a skull layer. Lines show the mean and shading the range across all one-at-a-time parameter variations; dashed lines mark the skull or brain boundaries. (b) Peak brain intensity when a single brain (gray) or skull (red) acoustic property is set to its LOW (open circles) or HIGH (filled circles) bound, in PRESTUS and BabelBrain. Bars span the two outcomes; dashed lines mark results with all properties at reference values. c, sound speed; ρ, density; α, attenuation coefficient; subscripts b and s denote brain and skull.

### 3.3 Skull properties differentially affect temperature rise in skull and at focus

Acoustic properties alone do not determine heating: temperature rise additionally depends on thermal tissue properties, and the amount of energy that is absorbed in the tissue. We next assessed the impact of acoustic and thermal parameter uncertainty on temperature rise in the skull and at the focus (Fig. 3). Skull absorption, the attenuated energy converted to heat, had the largest impact on temperature rise within the skull, followed by skull sound speed, density, attenuation, and specific heat capacity. Focal heating, relevant to the likelihood of thermal neuromodulation, depended on two sources of uncertainty. Skull properties determined how much energy reached the brain: higher skull attenuation reduced focal heating, with smaller contributions from skull density and sound speed (Fig. 3b). Brain absorption (selectively varied at fixed reference attenuation, and covarying with attenuation via the absorption fraction) determined how strongly energy heated brain tissue and was the largest single source of focal uncertainty both with and without a skull layer (Fig. 3c). All parameter effects were consistent across solvers.

**Figure 3.**
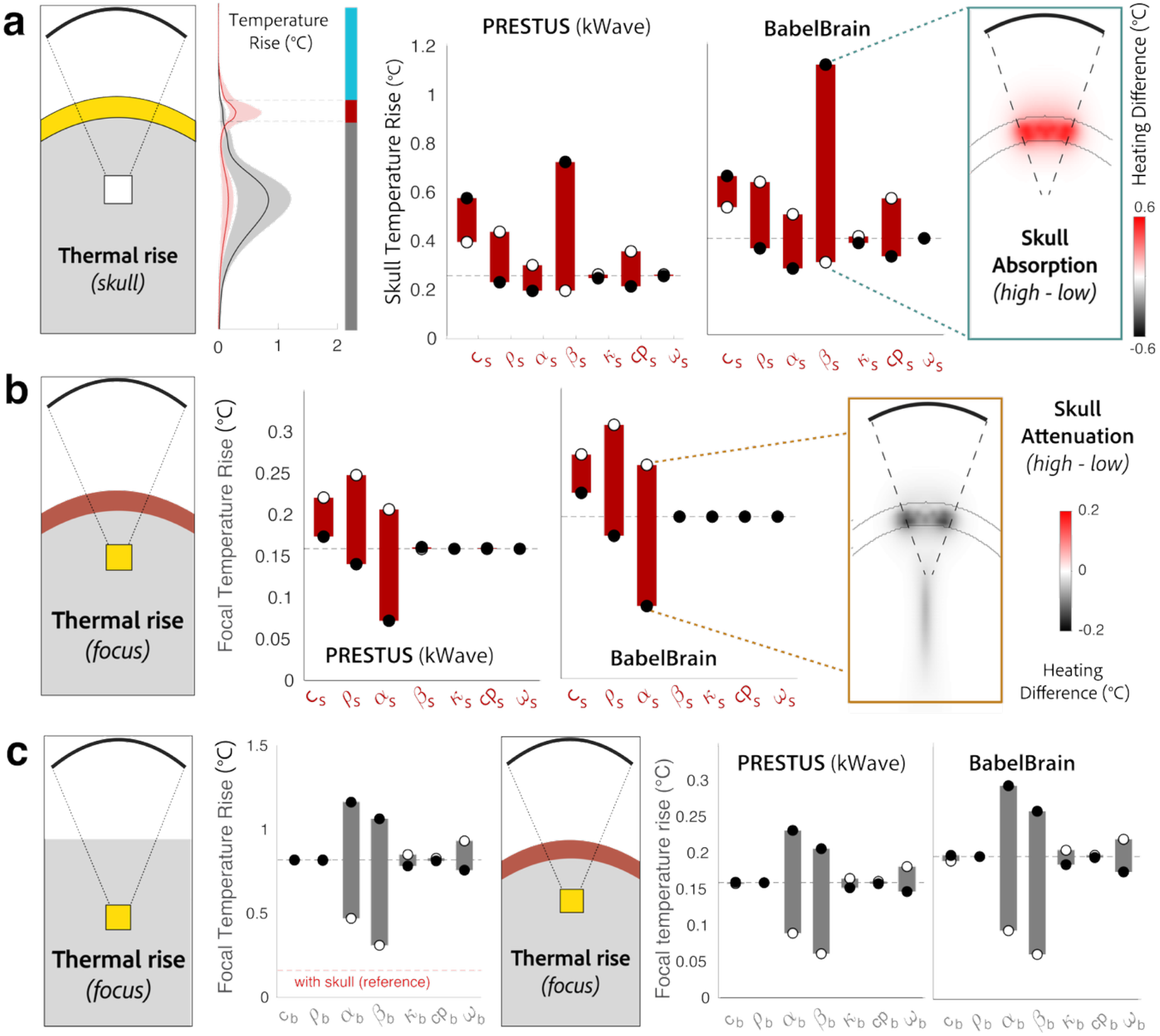
Parameter impact on temperature rise in the skull (a) and at the focus (b-c). Bars span peak temperature rise between LOW (open circles) and HIGH (filled circles) settings of each parameter, with all other parameters at reference values (dashed line). (a) Skull temperature rise for independent variation of skull parameters. Increased skull absorption has a large, but spatially limited impact on skull heating. (b) Focal temperature rise for independent variation of skull parameters. Larger skull attenuation reduces heating in both the skull layer and at the brain focus (at fixed absorption). Insets in a and b map the difference in peak temperature rise between the HIGH and LOW settings (based on PRESTUS results) of skull absorption (a) and skull attenuation (b) across the skull benchmark. Gray lines mark the inner and outer skull boundaries. (c) Focal temperature rise for variation of brain parameters without (left) and with (right) a skull layer. Brain absorption dominates focal uncertainty in both cases.

### 3.5 Generalizability of individual parameter impact to more transmissible skull

Skull thickness varies across regions and individuals, and transmission fluctuates non-linearly with thickness and frequency (Attali et al., 2023). To test whether the impact direction of acoustic and thermal parameters on exposure estimates observed with the 6.5 mm benchmark generalizes to other thicknesses, we repeated the sensitivity analysis for the most transmissive skull. A thickness sweep placed maximal transmission at 2.5 mm in our setup, while 6.5 mm lay on a falling flank between a local maximum (5.5 mm) and a local minimum (7.5 mm). The direction of all skull parameter effects was preserved, except for an inversion of the density effect on skull temperature rise specific to the most transmissive thickness (Supp. Fig. 2). Individual parameter effects were thus largely robust to skull thickness.

### 3.5 Uncertainty setups bound transmission and temperature rise of toolbox defaults

In practice, exposure uncertainty arises from the joint modeling of individually uncertain parameter estimates. To examine how far plausible property assumptions can move brain intensity and heating within the reported literature values, we combined parameter bounds into joint tissue-property configurations (Fig. 5). Temp-H and Temp-L combine, per property, the bounds that individually raised or lowered temperature rise in both skull and brain; Trans-H and Trans-L combine the acoustic bounds that raised or lowered transmission. We compared these joint setups to default property assumptions in three simulation platforms (Supp. Table S2). The toolboxes closely align on skull density and attenuation and yielded comparable brain intensity in the skull benchmark. Skull heating, however, varied markedly, in line with differences in skull absorption and heat capacity. k-Plan’s complete conversion of skull attenuation into absorption, which exceeds the upper literature bound, and its low heat capacity yielded the highest skull heating among the toolboxes. The soft tissue benchmark similarly showed variable heating at comparable intensity, reflecting differences in brain absorption (BabelBrain) and perfusion (none in k-Plan). These results suggest that diverging exposure estimates between toolboxes are expected given differing property assumptions; all toolbox estimates fell within the range spanned by the joint setups, although some default properties lie beyond the literature bounds.

## 4. Discussion

Uncertainty in tissue properties substantially reduces precision of ultrasound exposure estimates. Skull attenuation dominated uncertainty in brain intensity, skull absorption in skull heating, and brain absorption in focal heating. In line with prior reports (Aubry et al., 2022; Bouchoux et al., 2012; Krokhmal et al., 2025; Montanaro et al., 2021; Yoon et al., 2018), our results indicate a more limited impact of acoustic parameters on focal position and volume. The relative contribution of parameters to outcome uncertainty was robust across solvers, consistent with intensity converging within 10% between solvers sharing acoustic properties (Aubry et al., 2022). This highlights that solver differences alone cannot account for large discrepancies between k-Plan and BabelBrain in practice (Brandts et al., 2026). Our results instead explain the previous observation that peak intensity, but not temperature rise, correlated across platforms: platforms largely agree on attenuation-driven transmission loss (absent image-based property mapping), whereas thermal assumptions diverge.

Given the spread of reported tissue properties, simulated intracranial pressure and temperature rise should currently be interpreted as ranges rather than precise point estimates (Krokhmal et al., 2025; Stanziola et al., 2023). Explicit range estimation has already been proposed for image-based property mapping, in which uncertainty in the CT-to-sound-speed mapping alone propagates to up to ∼40 kPa standard deviation near the focus in a 60 kPa source-pressure simulation (Stanziola et al., 2023). Parameter setups proposed here provide a complementary, literature-based approach. Such setups remain valid across head geometries and protocols only if each property shifts outcomes in a fixed direction. This held for all properties except skull density, whose effect on skull heating inverted exclusively at the maximally transmissive thickness (Fig. 4; Supp. Fig. 2). Simulating multiple exposure assumptions can guard non-significant risk (NSR) evaluations against two systematic errors: overcompensation of output due to underestimated exposure, and underdosing due to overestimated temperature rise. We recommend that simulation tools and their users routinely report tissue property choices (Martin et al., 2024), provide free-field parameters alongside model-based estimates, and disclose uncertainty around point estimates where possible.

**Figure 4.**
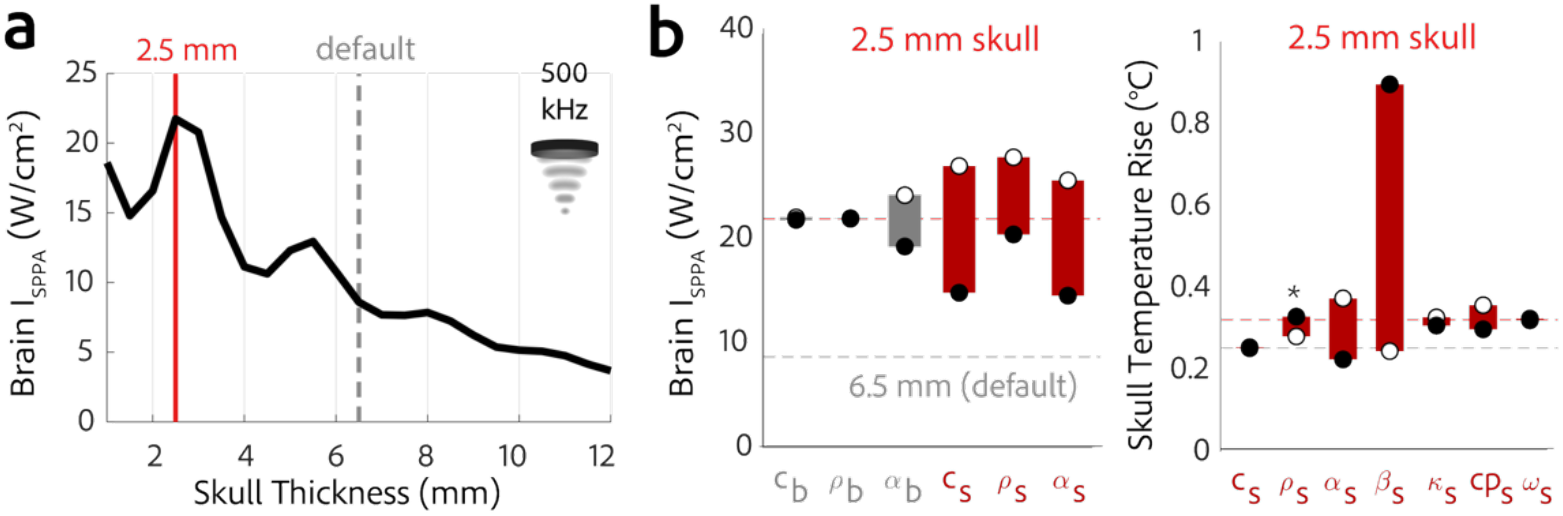
Impact of bone transmission on parameter impact. (a) Transmission at the fundamental frequency varies as a function of skull thickness (cf. Attali et al., 2023) due to resonance effects. (b) Skull parameter impact on estimated in-situ intensity and skull temperature rise for the max. transmissive skull benchmark. An inversion in effect direction of density on skull heating is indicated via * (cf. Fig. 3a) but was constrained to the most transmissive skull thickness (see Supp. Fig. 2).

Our sensitivity analysis indicates that exposure uncertainty arises primarily from limited and variable measurements of skull attenuation and skull/brain absorption. A previous sensitivity analysis instead ranked sound speed as the most sensitive skull property for peak pressure (Robertson, Martin, et al., 2017). The two results are compatible: Robertson et al. perturbed properties relative to reference values and found that keeping peak-pressure errors below 5% required sound speed to stay within 4% of its reference across 0.5–1 MHz, whereas attenuation could deviate by up to 20% at 500 kHz. The literature range for attenuation spans a factor of 3, far exceeding that tolerance. Per unit change, sound speed may matter most but across the empirical range, attenuation and absorption dominate exposure uncertainty. Similarly, Leung et al. (2019) demonstrated that skull attenuation across a plausible range (1–2 Np/cm at 680 kHz; ∼12.8–25.5 dB/cm/MHz) produced an approximately four-fold range in simulated focal temperature rise (cf. Fig. 3b). Our results identify brain absorption, for which no human measurements were identified, as a major contributor.

Skull attenuation combines scattering, which reduces transmission, and absorption, the conversion of acoustic into thermal energy (Liu et al., 2019). Toolboxes have frequently coupled the two via an absorption fraction applied during the thermal estimation step, with defaults from 16.3% (BabelBrain) to 100% (k-Plan) predicting strongly divergent skull heating (Fig. 5). Fractions tie heat deposition to a wide, scattering-dominated attenuation range, and because scattering rises more steeply with frequency than absorption, fractions derived near 1 MHz likely underestimate the absorbed share at typical TUS frequencies (Liu et al., 2019). We therefore propose to specify skull absorption as a dedicated coefficient. However, few measurements separate the two: Pinton et al. (2012) estimated a longitudinal absorption coefficient of 2.7 dB/cm at 1 MHz in ex vivo human cortical bone, consistent with 2.8 dB/cm/MHz across 1–10 MHz in formalin-stored bovine compact bone (D. N. White et al., 1978). No measurements exist for the diploë. Because roughly half of its volume is marrow and its losses are dominated by scattering, its absorption per unit length is presumedly at or below that of cortical bone, such that cortical bounds bracket or exceed the absorption of a homogenized skull layer.

**Figure 5.**
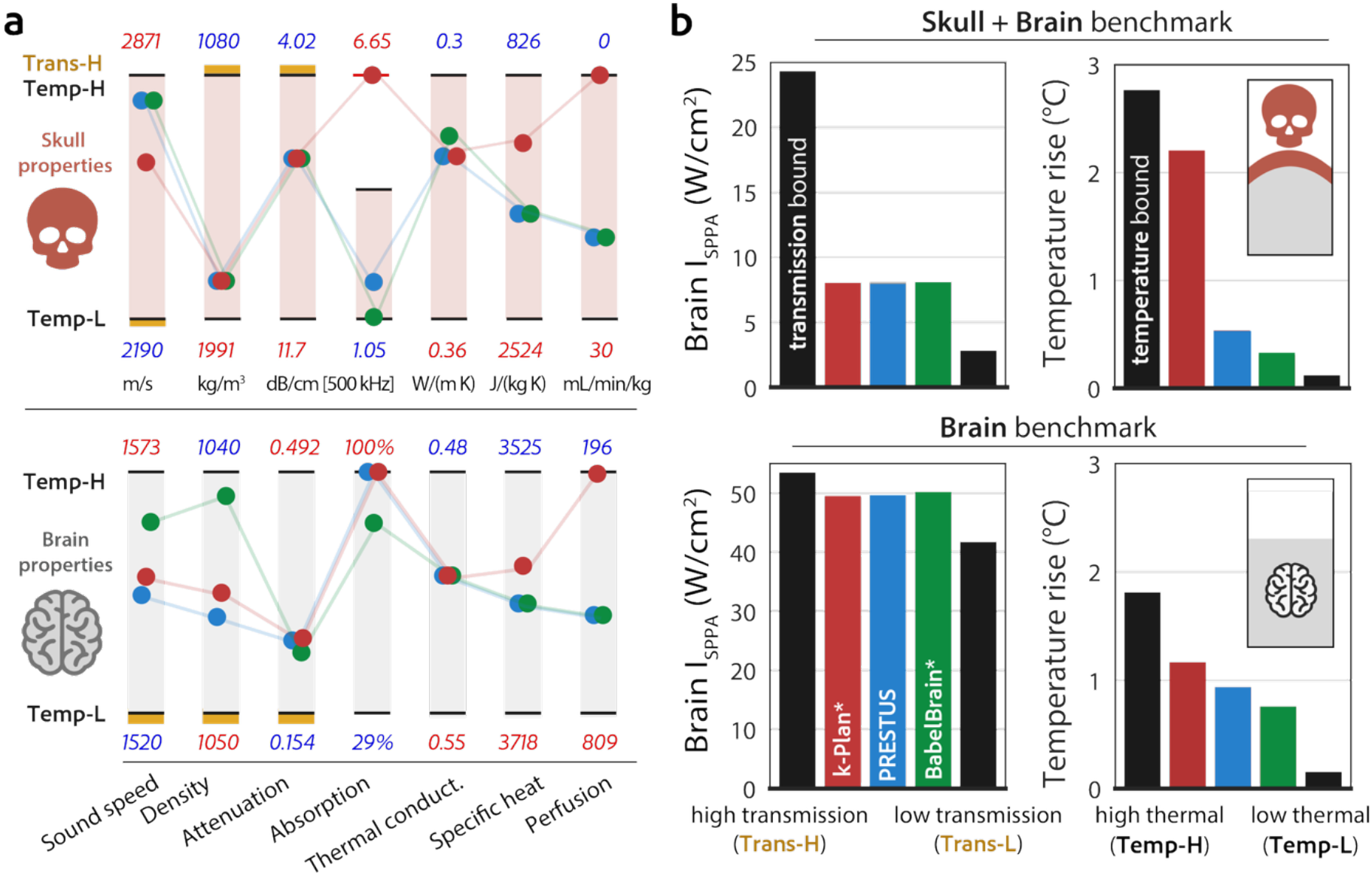
Joint parameter bounds for transmission and global temperature rise. (a) Joint parameter setups of skull and brain tissue properties and comparison to toolbox defaults (see Supp. Table S2). Temperature (Temp-H) represents, per property, the literature bound that increased skull and brain temperature rise in the one-at-a-time sensitivity analysis, while Temp-L represents the opposite. The shaded area for each parameter represents its modeled property range. Values above and below indicate the direction of parameter settings for the Temp-H and -L settings. For skull absorption, this range has an upper bound by the k-Plan value. High transmission (Trans-H) represents, per acoustic property, the bound that increased transmission to the brain, and Trans-L the parameter bound that reduced it. Trans-H bounds are shown in yellow; Trans-L represents the opposite bound (not shown). (b) I_SPPA_ and global temperature rise estimates for the parameter bounds and the toolbox defaults in the skull + brain benchmark (top) and the brain benchmark (bottom). The skull benchmark jointly models brain and skull tissue bounds. * Both k-Plan and BabelBrain parameters were emulated in the k-Wave backend of PRESTUS and do not capture other toolbox differences in parameter mapping and simulation workflow.

Compressional-only solvers may systematically underestimate skull heating and misestimate transmission when shear mode conversion is present. At the skull surface, part of the incident compressional energy converts into shear waves, particularly at oblique incidence (Bouchoux et al., 2012; Gao et al., 2026; Pichardo et al., 2017; Pinton et al., 2012; P. J. White et al., 2006). Shear waves accounted for on average 21% of skull temperature rise in a monkey skull model (Wang et al., 2018), and shear heating increased sharply beyond 20–30° incidence at 0.8–1.2 MHz in a human skull model (Kilinc et al., 2025). For conservative simulations of skull heating, we recommend that compressional-only solvers use a higher absorption coefficient than our reference. Neglecting shear waves also biases transmission estimates. A fluid model overestimated intracranial pressure in an ex vivo human skull at 500 kHz by on average 40%, compared with 20% for an elastic model (Gao et al., 2026). Both models, however, assumed a longitudinal attenuation of 4.34 dB/cm at 500 kHz (P. J. White et al., 2006), close to our LOW bound, whereas a higher attenuation within the uncertainty range may reduce this discrepancy. More generally, a single scalar parameter cannot model mode conversion which varies with incidence angle, skull curvature, and microstructure. Simulations that resolve skull microstructure, ideally from high-resolution micro-CT (Robertson, Cox, et al., 2017) promise to overcome these limitations toward improved estimates of both transmission and heating.

Our simulations involve simplifying assumptions. First, we modeled a homogeneous skull layer in an axisymmetric geometry. Compartment-resolved attenuation measurements that could inform multi-layer models exist for only a handful of specimens (Webb et al., 2021), with fewest datapoints at neuromodulation-relevant frequencies (Supp. Fig. 3), where longer wavelengths complicate layer-resolved measurement. The benchmark further omits scalp, hair, meninges and CSF, treats brain tissue as uniform, contains no contralateral skull, and assumes spatially uniform perfusion. These simplifications were made for efficient numerical study of the sensitivity to the key parameters identified using earlier more detailed models. Second, we treated parameters as independent, whereas in tissues they are jointly determined by microstructure (mineral content, porosity, and water fraction). In human soft tissue, sound speed and density are strongly correlated, whereas attenuation shows no robust relation to either (Mast, 2000). In cranial bone, the interrelation of sound speed, attenuation, and porosity at the level of skull microstructure forms the physical basis for density- or CT-based acoustic property mapping (Marsac et al., 2017; Pichardo et al., 2011; Webb et al., 2018), and for microstructure–insertion loss relations (Riis et al., 2022). Comparisons of well-controlled skull experiments with simulations can thus indicate smaller discrepancies than uncertainty across the full and methodologically variable literature range would predict.

Our analysis identifies where measurement and modeling effort would most reduce exposure uncertainty. Image-based property maps derived from (micro-)CT can capture patient-specific spatial variation in porosity and density, but accurate mappings require density phantoms, relations remain variable for sound speed (Clinard et al., 2025b; Montanaro et al., 2021; Stanziola et al., 2023) and especially attenuation (Clinard et al., 2025a; Leung et al., 2019; Webb et al., 2021), and MRI-based surrogates add further mapping uncertainty (Miscouridou et al., 2022; Yaakub et al., 2023). Given the large variation in attenuation measured across multiple studies and methodologies, this uncertainty in acoustic parameter mapping from clinical images may be a real feature of skull acoustic properties (Clinard et al., 2025a, 2025b). One way to mitigate this uncertainty is to use subject-specific transmission measurements (Dadgar-Kiani et al., 2026; Riis et al., 2024). These measurements can directly estimate focal intensity or serve as an additional constraint on simulation. Such calibrations measure total insertion loss and therefore cannot constrain skull absorption, which dominated uncertainty in skull, or brain absorption which strongly impacted focal heating. Narrowing this uncertainty requires new measurements: we found no measurements of human brain absorption and only two skull absorption measurements, one of them in human tissue and both in cortical bone alone. Dedicated measurements on multiple human ex-vivo skulls spanning realistic thickness and porosity, across 0.2–1 MHz, quantifying absorption and attenuation on the same specimens and separating them for cortical and diploic bone, together with measurements of human brain absorption, would address the largest source of uncertainty in thermal safety estimation. The ITRUSST Tissue Properties Database (Kosciessa, Webb, et al., 2026) provides a venue to register new measurements once they become available.

Beyond tissue properties, simulation frameworks face further uncertainty sources (Angla et al., 2023): for example, multi-element transducer specification (McDannold et al., 2019; Pasquinelli et al., 2020) and tissue orientation in the simulation domain both affect predicted focal intensity (Attali et al., 2023; Robertson, Cox, et al., 2017). The methodology established here extends to these sources, to identify which dominate simulation outcomes and should be prioritized toward robust personalized ultrasound simulations.

## CRediT authorship contribution statement

Julian Q. Kosciessa: Conceptualization, Methodology, Software, Formal analysis, Visualization, Writing – original draft, Writing – review & editing.

Taylor Webb: Conceptualization, Methodology, Data curation, Writing – review & editing.

Elly Martin: Conceptualization, Data curation, Writing – review & editing.

Brad Treeby: Conceptualization, Methodology, Writing – review & editing.

Sanjay Yengul: Conceptualization, Writing – review & editing.

Lennart Verhagen: Conceptualization, Methodology, Supervision, Writing – review & editing.

Kim Butts Pauly: Conceptualization, Writing – review & editing, Supervision.

Samuel Pichardo: Conceptualization, Methodology, Software, Formal analysis, Visualization, Resources, Writing – review & editing.

## Acknowledgments

This work was supported by a Radboud Excellence Initiative grant to JQK and a VIDI Fellowship by the Dutch Research Council (NWO 18919) to LV. We thank all members of the ITRUSST acoustic properties working group for their assistance in compiling and discussing the property literature. We thank Sanjay Yengul for his contributions to conceptualization, discussion, review, and editing.

