## Supplemental Materials for "Impact of acoustic property uncertainty on transcranial ultrasound simulation"

### Supplementary Materials

**Supplementary Table S1.** Literature-derived acoustic and thermal property values for skull (S1a), brain (S1b) and water (S1c). Within each property, values are listed in ascending order. Attenuation and absorption are converted to dB/cm (1 Np = 8.686 dB) and scaled to 0.5 MHz with the reported or study-fitted power-law exponent b; "(assumed)" marks b = 1 where no exponent was reported. Test frequency gives the measurement frequency or range; where a value was fitted across several discrete frequencies, the closest to 500 kHz is used. Blue and red rows set the LOW and HIGH bounds of Table 1. Gray rows were not used for the bounds; the reason is given in the Reference column or a footnote. "A: B" in the Reference column denotes a value compiled in A from source(s) B. Bone values from non-cranial sites (e.g. femur, vertebrae) or non-human species are listed under skull as proxies and labeled by site and species.

#### Supplementary Table S1a. Skull

| SOUND SPEED (m/s) | | | |
|---|---|---|---|
| **Tissue** | **Value** | **Test frequency** | **Reference** |
| Skull (cancellous) | 2190 m/s | 0.3–0.8 MHz | (Fry & Barger, 1978) |
| Skull (cancellous) | 2240 m/s | 0.3–0.8 MHz | (Fry & Barger, 1978) |
| Skull (whole) | 2291 m/s | 0.27 MHz | (Pichardo et al., 2011) |
| Skull (whole) | 2337 m/s | 0.3–0.8 MHz | (Fry & Barger, 1978) |
| Skull (whole) | 2408 m/s | 0.836 MHz | (Pichardo et al., 2011) |
| Skull (parietal) | 2486 m/s | 0.5 MHz | (Riis et al., 2022) |
| Skull (temporal) | 2541 m/s | 1.03 MHz | (Ammi et al., 2008) |
| Skull (occipital) | 2603 m/s | 0.5 MHz | (Riis et al., 2022) |
| Skull (whole) | 2609 m/s | 0.5 MHz | (Riis et al., 2022) |
| Skull (whole) | 2740 m/s | 0.3–0.8 MHz | (Fry & Barger, 1978) |
| Skull (whole) | 2820 m/s | 1 MHz | (P. J. White et al., 2006) |
| Skull (frontal) | 2871 m/s | 0.5 MHz | (Riis et al., 2022) |

| DENSITY (kg/m³) | | |
|---|---|---|
| **Tissue** | **Value** | **Reference** |
| Skull (cancellous; human vertebrae/calcaneus, marrow intact) | 1080 kg/m³ | (Blanton & Biggs, 1968) |
| Skull (cancellous; human) | 1105 kg/m³ | (Clauser et al., 1969) |
| Skull (cancellous; dog) | 1350 kg/m³ | (R. A. Robinson, 1958) |
| Skull (cortical; human) | 1800 kg/m³ | (Clauser et al., 1969) |
| Skull (cortical; dog) | 1800 kg/m³ | (R. A. Robinson, 1958) |
| Skull (whole) | 1840 kg/m³ | (Fry & Barger, 1978; via Pichardo et al., 2011) |
| Skull (cortical; human tibia/femur/humerus) | 1850 kg/m³ | (Blanton & Biggs, 1968) |
| Skull (whole) | 1904 kg/m³ | (Pichardo et al., 2011) |
| Skull (cortical; human femur/tibia, marrow-free) | 1991 kg/m³ | (Gong et al., 1964) |
| Skull (cortical; unspecified) | 2100 kg/m³ | (Henschel, 1943) [1] |

[1] Single measurement on fresh cortical bone of unspecified species and anatomical site in a study of dentin heating (Henschel, 1943, p. 325).

| LONGITUDINAL ATTENUATION COEFFICIENT (dB/cm @0.5MHz) | | | | | | |
|---|---|---|---|---|---|---|
| **Tissue** | **Reported value** (units vary) | **Test frequency** | **Exponent b** | **dB/cm @ 1 MHz** | **dB/cm @ 0.5 MHz** | **Reference** |
| Skull (cancellous) | $\alpha_0$ = 47 Np/m | 1 MHz | 1.2 | 4.08 dB/cm | 1.78 dB/cm | (Baumgartner et al., 2025): (El-Brawany et al., 2009) [2] |
| Skull (cortical) | 54.553 Np/m/MHz | 1 MHz | 1 | 4.74 dB/cm | 2.37 dB/cm | (Baumgartner et al., 2025): (Goss et al., 1980; Hosokawa & Otani, 1997) [3] |
| Skull (whole) | 53 Np/m | 0.548 MHz | 1.47 (fitted) | 11.1 dB/cm | 4.02 dB/cm | (P. J. White et al., 2006) [4] |
| Skull (temporal; effective, model-calibrated) | 15.3 dB/cm/MHz | 0.5 MHz | 1.5 | 15.3 dB/cm | 5.41 dB/cm | (Dadgar-Kiani et al., 2026) |
| Skull (outer table) | 0.7 Np/cm | 0.5 MHz | 1.3 | 15.0 dB/cm | 6.1 dB/cm | (Webb et al., 2021) |
| Skull (whole) | 33.8 Np/m | 0.27 MHz | 1.44 | 20.8 dB/cm | 7.65 dB/cm | (Pichardo et al., 2011) [5] |
| Skull (whole) | 16.6 dB/cm | 1 MHz | 1.00 (assumed) | 16.6 dB/cm | 8.3 dB/cm | (Pinton et al., 2012) [6] |
| Skull (outer table) | 1.67 Np/cm | 1 MHz | 0.79 | 14.51 dB/cm | 8.38 dB/cm | (Fry & Barger, 1978) |
| Skull (temporal) | 154–713 Np/m | 1.03 MHz | 1.19 | 21.3 dB/cm | 9.35 dB/cm | (Ammi et al., 2008) [7] |
| Skull (inner table) | 2.15 Np/cm | 1 MHz | 0.90 | 18.67 dB/cm | 10.01 dB/cm | (Fry & Barger, 1978) |
| Skull (inner table) | 2.9 Np/cm | 1 MHz | 1.3 | 25.2 dB/cm | 10.2 dB/cm | (Webb et al., 2021) |
| Skull (diploë) | 1.2 Np/cm | 0.5 MHz | 1.3 | 25.7 dB/cm | 10.4 dB/cm | (Webb et al., 2021) |
| Skull (whole) | 2.48 Np/cm | 1 MHz | 1 | 21.54 dB/cm | 10.77 dB/cm | (Fry & Barger, 1978) |
| Skull (diploë) | 3.68 Np/cm | 1 MHz | 1.45 | 31.98 dB/cm | 11.70 dB/cm | (Fry & Barger, 1978) [8] |
| Skull (whole) | 176 Np/m | 0.5 MHz | 1 | 30.57 dB/cm | 15.29 dB/cm | (Riis et al., 2022) [9] |
| Skull (whole) | 208 Np/m | 0.5 MHz | 1 | 36.13 dB/cm | 18.07 dB/cm | (Riis et al., 2022) |
| Skull (whole) | 489 Np/m | 0.5 MHz | 1 | 84.95 dB/cm | 42.47 dB/cm | (Riis et al., 2022) |

| LONGITUDINAL ABSORPTION COEFFICIENT (dB/cm @0.5MHz) | | | | | | |
|---|---|---|---|---|---|---|
| **Tissue** | **Reported value** (units vary) | **Test frequency** | **Exponent b** | **dB/cm @ 1 MHz** | **dB/cm @ 0.5 MHz** | **Reference** |
| Skull (cortical; bovine ivory bone, fresh) | 2.1 dB/cm/MHz | 1.5–8.0 MHz | 1.00 | 2.1 dB/cm | 1.05 dB/cm | (D. N. White et al., 1978) sample 3 (4.5 mm biplanar slice; fresh bone); |
| Skull (cortical; longitudinal wave) | 2.7 dB/cm | 1 MHz | 1.00 (assumed) | 2.7 dB/cm | 1.35 dB/cm | (Pinton et al., 2012) |
| Skull (cortical; bovine ivory bone, formalin-stored) | 2.6 dB/cm/MHz | 1.95–7.25 MHz | 1.00 | 2.6 dB/cm | 1.30 dB/cm | (D. N. White et al., 1978) sample 2 (4.2 mm biplanar slice; same bone as sample 1) |
| Skull (cortical; bovine ivory bone, formalin-stored) | 2.8 dB/cm/MHz | 1–10 MHz | 1.00 | 2.8 dB/cm | 1.40 dB/cm | (D. N. White et al., 1978) sample 1 (4.9 mm biplanar slice; formalin-stored) |
| Skull (whole attenuation) | 53 Np/m | 0.548 MHz | 1.47 | 11.1 dB/cm | 4.02 dB/cm | (P. J. White et al., 2006): low end of attenuation = proxy for high end of absorption |

---

[2] El-Brawany et al. (2009): porcine bone, n = 4 samples, bone type (cortical/cancellous) not specified; 5 MHz broadband pulse-echo (−6 dB bandwidth) at 37 °C. The paper reports b = 1.37 (IT'IS lists 1.2). The pig scapula and sheep skull samples were used for thermal measurements only.

[3] Bovine femur (cow thigh bones), 0.5–5 MHz range. Hosokawa & Otani (1997) studied cancellous bone; cortical values are given in their appendix. Goss et al., 1980 meta-analysis includes most values measured at > 1MHz.

[4] White et al. (2006, Table 3) report longitudinal attenuation of 14 ± 17, 53 ± 43 and 70 ± 28 Np/m at 0.272, 0.548 and 0.840 MHz. The value at 0.5 MHz was scaled from the measurement nearest the target frequency: 53 Np/m = 4.60 dB/cm at 0.548 MHz, and 4.60 × (0.5/0.548)^1.47 = 4.02 dB/cm, where b = 1.47 is the least-squares power-law exponent across the three frequencies. Evaluating the fitted power law itself at 0.5 MHz would instead give 3.24 dB/cm. Relative to the measured 4.60 dB/cm at 0.548 MHz, 3.24 dB/cm is consistent with this measurement only for an unrealistically high local exponent of ln(4.60/3.24) / ln(0.548/0.5) ≈ 3.8.

[5] Value evaluated from the study's power-law fit across the reported frequencies.

[6] 16.6 dB/cm at 1 MHz is derived from insertion loss through skull samples, so it is not a reflection-corrected attenuation.

[7] Ammi et al. (2008, Table 5) report temporal-bone attenuation for five skulls at 1.03 MHz (154–713 Np/m) and 2.00 MHz (379–1210 Np/m). The value at 0.5 MHz was scaled from the measurements nearest the target frequency: the geometric mean across skulls at 1.03 MHz, 254 Np/m = 22.1 dB/cm, gives 22.1 × (0.5/1.03)^1.19 = 9.35 dB/cm, where b = 1.19 is the least-squares power-law exponent across all ten values. Reflection loss was corrected at the front surface only, the samples were dried temporal bone, and b was fitted above the target frequency (1.03–2.00 MHz).

[8] Attenuation coefficients for diploë and whole skull are not tabulated in Fry & Barger (1978); values were digitized from Fig. 10 (diploë, skull samples No. 6 and No. 8) and Fig. 11 (full skull No. 8) respectively.

[9] These outlier attenuation values were derived from a single-term model, T = R·exp(−αh), where α is the slope of log pressure transmission against thickness across a skull's segments, and the reflection loss R is a free intercept that is fitted, not measured or reported. This makes the questionable assumption that reflection loss and bone composition stay constant as thickness changes. Phase cancellation at the large unfocused receiver was also not corrected. The resulting coefficients varied almost three-fold across the three skulls, and the authors describe the model as "likely simplistic".

| THERMAL CONDUCTIVITY (W/m/K) | | |
|---|---|---|
| Tissue | Value | Reference |
| Skull (cancellous; cow femur) | 0.287 W/m/K | (Clattenburg et al., 1975); non-adult sample |
| Skull (cortical; human femur) | 0.300 W/m/K | (McIntosh & Anderson, 2010) |
| Skull (cancellous; human femur) | 0.300 W/m/K | (McIntosh & Anderson, 2010) |
| Skull (cancellous) | 0.3125 W/m/K | (Baumgartner et al., 2025):<br>(Collins et al., 2004; McIntosh & Anderson, 2010) |
| Skull (cortical) | 0.32 W/m/K | (Baumgartner et al., 2025):<br>(Clattenburg et al., 1975; Collins et al., 2004; McIntosh & Anderson, 2010) |
| Skull (cortical) | 0.36 W/m/K | (Collins et al., 2004) |
| Skull (cancellous) | 0.36 W/m/K | (Collins et al., 2004) |

| SPECIFIC HEAT CAPACITY (J/kg/K) | | |
|---|---|---|
| Tissue | Value | Reference |
| Skull (cortical, min; pig & sheep) | 826 J/kg/K | (McIntosh & Anderson, 2010) |
| Skull (cortical; human) | 1255 J/kg/K | (McIntosh & Anderson, 2010) |
| Skull (cortical; human) | 1300 J/kg/K | (Duck, 1990) |
| Skull (cortical) | 1300 J/kg/K | (Collins et al., 2004) |
| Skull (cortical) | 1313 J/kg/K | (Baumgartner et al., 2025):<br>(Collins et al., 2004; Duck, 1990; McIntosh & Anderson, 2010) |
| Skull (cortical; cow) | 1330 J/kg/K | (Duck, 1990) |
| Skull (cortical; pig) | 1360 J/kg/K | (Duck, 1990) |
| Skull (cortical; human femur) | 1650 J/kg/K | (McIntosh & Anderson, 2010) |
| Skull (cancellous; human femur) | 2060 J/kg/K | (McIntosh & Anderson, 2010) |
| Skull (cancellous) | 2238 J/kg/K | (Collins et al., 2004) |
| Skull (cancellous) | 2274 J/kg/K | (Baumgartner et al., 2025):<br>(Collins et al., 2004; McIntosh & Anderson, 2010) |
| Skull (cancellous; cow femur) | 2524 J/kg/K | (McIntosh & Anderson, 2010):<br>(Clattenburg et al., 1975) |

| PERFUSION (mL/min/kg) | | |
|---|---|---|
| Tissue | Value | Reference |
| Skull (cortical; assumed, not measured) | 0 mL/min/kg | (McIntosh & Anderson, 2010) |
| Skull (cancellous; human, bone site not stated) | 12 mL/min/kg | (McIntosh & Anderson, 2010) |
| Skull (cortical) | 14 mL/min/kg | (Collins et al., 2004) |
| Skull (cancellous) | 30 mL/min/kg | (Collins et al., 2004) |

### Supplementary Table S1b. Brain

| SOUND SPEED (m/s) | | | |
|---|---|---|---|
| Tissue | Temperature | Value | Reference |
| Brain (gray matter) | - | 1500 m/s | (Baumgartner et al., 2025): noted as arbitrary |
| Brain (gray matter; sheep) | ~ 20°C | 1520 m/s | (Labuda et al., 2022) |
| Brain (human) | 22 °C | 1532 m/s | (Duck, 1990) |
| Brain (global, sheep) | ~20 °C | 1535 m/s | (Labuda et al., 2022) |
| Brain (white matter, sheep) | ~20 °C | 1537 m/s | (Labuda et al., 2022) |
| Brain (white matter) | 22/40 °C | 1552.5 m/s | (Baumgartner et al., 2025): (Duck, 1990) |
| Brain (global, human) | 37 °C | 1562–1567 m/s | (Kremkau et al., 1981) |
| Brain (white matter, human; corona radiata) | 37 °C | 1568 m/s (interpolated) | (Kremkau et al., 1981) |
| Brain (white matter, human; corona radiata) | 40 °C | 1573 m/s | (Kremkau et al., 1981, via Duck, 1990) |

| DENSITY (kg/m³) | | |
|---|---|---|
| Tissue | Value | Reference |
| Brain (white matter; human) | 1040 kg/m³ | (Cooper & Trezek, 1972) |
| Brain (white matter) | 1040 kg/m³ | (Huang & Wu, 1976) |
| Brain (cow) | 1041 kg/m³ | (Poppendiek et al., 1966) |
| Brain (white matter) | 1041 kg/m³ | (Baumgartner et al., 2025): (Cooper & Trezek, 1972; Danilewsky, 1863; Huang & Wu, 1976) |
| Brain (gray matter) | 1044.5 kg/m³ | (Baumgartner et al., 2025): (Cooper & Trezek, 1972; Danilewsky, 1863) |
| Brain | 1045.5 kg/m³ | (Baumgartner et al., 2025): (Cooper & Trezek, 1972; Poppendiek et al., 1966) |
| Brain (gray matter) | 1050 kg/m³ | (Cooper & Trezek, 1972) |

| LONGITUDINAL ATTENUATION COEFFICIENT (dB/cm @0.5MHz) | | | | | | |
|---|---|---|---|---|---|---|
| **Tissue** | **Reported value** (units vary) | **Test frequency** | **Exponent b** | **dB/cm @ 1 MHz** | **dB/cm @ 0.5 MHz** | **Reference** |
| Brain (gray matter) | 1.2 Np/m/MHz$^b$ | - | 1.21 | 0.104 dB/cm | 0.045 dB/cm | (Baumgartner et al., 2025); no source; matches GM absorption in (Lyons & Parker, 1988) |
| Brain (cow, white/gray mix; 20–22 °C) | 0.040 Np/cm (@ 1 MHz) | 1–13 MHz | 1.17 | 0.347 dB/cm | 0.154 dB/cm | (Lyons & Parker, 1988) |
| Brain (human) | 0.435 dB/cm/MHz$^b$ | 1–5 MHz | 1.08 | 0.435 dB/cm | 0.206 dB/cm | (Duck, 1990) [10] |
| Brain (human, mixed gray/white) | 0.58 dB/cm/MHz$^b$ | 1–6 MHz | 1.30 | 0.580 dB/cm | 0.236 dB/cm | (Bamber, 1981) |
| Brain (white matter) | 6.7082 Np/m/MHz$^b$ | - | 1.09 | 0.583 dB/cm | 0.274 dB/cm | (Baumgartner et al., 2025): (Duck, 1990) [11] |
| Brain (sheep, fixed; lowest spatial mean) | 0.59 dB/cm/MHz$^b$ | 3.5–10 MHz | 1.00 | 0.590 dB/cm | 0.295 dB/cm | (Labuda et al., 2022) |
| Brain (cow, cerebral cortex; 25 °C) | 0.14 Np/cm | 0.7–5 MHz | 1.05 | 0.697 dB/cm | 0.336 dB/cm | (Etoh et al., 1994)[12] |
| Brain (human, white matter) | 0.8 dB/cm/MHz | 1–6 MHz | 1.10 | 0.800 dB/cm | 0.373 dB/cm | (Bamber, 1981) |
| Brain (human adult; summary trend) | 0.84 dB/cm/MHz$^b$ | ~1–10 MHz | 1.10 | 0.840 dB/cm | 0.392 dB/cm | (Bamber, 1998) |
| Brain (cat, in vivo; 37 °C) | 0.28–0.50 Np/cm | 4.2 MHz | 1.00 (assumed) | 0.807 dB/cm | 0.403 dB/cm | (T. C. Robinson & Lele, 1972)[13] |
| Brain (human, gray matter) | 0.78 dB/cm/MHz$^b$ | 1–5 MHz | 0.95 | 0.78 dB/cm | 0.404 dB/cm | (Kremkau et al., 1981) |
| Brain (sheep, fixed; highest spatial mean) | 0.81 dB/cm/MHz$^b$ | 3.5–10 MHz | 1.00 | 0.810 dB/cm | 0.405 dB/cm | (Labuda et al., 2022) |
| Brain (human, all sites) | 0.87 dB/cm/MHz$^b$ | 1–5 MHz | 1.08 | 0.870 dB/cm | 0.412 dB/cm | (Kremkau et al., 1981) |
| Brain (human, mixed: basal ganglia) | 0.90 dB/cm/MHz$^b$ | 1–5 MHz | 0.94 | 0.900 dB/cm | 0.469 dB/cm | (Kremkau et al., 1981) |
| Brain (human, white matter) | 1.04 dB/cm/MHz$^b$ | 1–5 MHz | 1.08 | 1.04 dB/cm | 0.492 dB/cm | (Kremkau et al., 1981) |

| LONGITUDINAL ABSORPTION COEFFICIENT (dB/cm @0.5MHz) [14] | | | | | | |
|---|---|---|---|---|---|---|
| **Tissue** | **Reported value** (units vary) | **Test frequency** | **Exponent b** | **dB/cm @ 1 MHz** | **dB/cm @ 0.5 MHz** | **Reference** |
| Brain (cow; gray matter; 20–22 °C) | 0.012 Np/cm | 1–13 MHz | 1.21 | 0.104 dB/cm | 0.045 dB/cm | (Lyons & Parker, 1988) |
| Brain (compiled; non-human animal data, 37 °C) | 10–28 dB/m | 1 MHz | 1.00 (assumed) | 0.10–0.28 dB/cm | 0.05–0.14 dB/cm | (International Commission on Radiation Units and Measurements, 1998) |
| Brain (cat) | 0.12 dB/cm | 0.7 MHz | 1.18 | 0.183 dB/cm | 0.081 dB/cm | (Goss et al., 1979) |
| Brain (cat; power-law fit) | 0.21 dB/cm/MHz | 0.7–7 MHz | 1.18 | 0.210 dB/cm | 0.093 dB/cm | (Goss et al., 1979) |
| Brain (cow; white matter; 20–22 °C) | 0.064 Np/cm | 1–13 MHz | 1.27 | 0.556 dB/cm | 0.231 dB/cm | (Lyons & Parker, 1988) |
| Brain (upper limit) | - | - | - | - | ≤ attenuation | conservative limit |

| THERMAL CONDUCTIVITY (W/m/K) | | |
|---|---|---|
| **Tissue** | **Value** | **Reference** |
| Brain (white matter) | 0.48 W/m/K | (McIntosh & Anderson, 2010) |
| Brain | 0.51 W/m/K | (McIntosh & Anderson, 2010) |
| Brain (gray matter) | 0.55 W/m/K | (McIntosh & Anderson, 2010) |

[10] Duck (1990, Table 4.16) lists this entry as human white matter, citing Kremkau et al. (1981). It equals half the value Kremkau et al. report for all adult brain samples (0.87 dB/cm at 1 MHz; identical exponent, temperature, and frequency range).

[11] The IT'IS V5.0 white-matter attenuation (6.7082 Np/m/MHz, b = 1.09) equals the geometric mean of two entries in Duck (1990, Table 4.16): Bamber's (1981) white-matter value as rounded by Duck (0.09 Np/cm; 0.8 dB/cm in Bamber, 1981) and the adjacent entry for human brain from Kremkau et al. (1981), which Duck lists at half its original value (0.050 Np/cm); b is the arithmetic mean of their exponents (1.1, 1.08). With the original Kremkau value (0.100 Np/cm), the geometric mean would be 9.49 Np/m/MHz (0.387 dB/cm at 0.5 MHz).

[12] Reported as an absorption coefficient (resonance method); listed as attenuation because the resonance width captures the total loss in the sample. The reported unit ($cm^{-1}$) is read as Np/cm; read as dB/cm, the value would be 0.039 dB/cm at 0.5 MHz, an order of magnitude below the other brain values.

[13] Robinson & Lele (1972, p. 1345) report that most attenuation coefficients across 20 cats fell between 0.28 and 0.50 Np/cm at 4.2 MHz; no mean or individual values are given. The tabulated value is the midpoint of this range, scaled with b = 1, which matches the authors' own linear scaling to 2.7 MHz (Table I).

[14] No human brain absorption measurements were identified in the reviewed literature.

| SPECIFIC HEAT (J/kg/K) | | |
|---|---|---|
| **Tissue** | **Value** | **Reference** |
| Brain (white matter; human) | 3525 J/kg/K | (McIntosh & Anderson, 2010) |
| Brain (white matter) | 3582.8 J/kg/K | (Baumgartner et al., 2025):<br>(Bernardi et al., 2003; Duck, 1990; McIntosh & Anderson, 2010) |
| Brain (white matter; human) | 3600 J/kg/K | (Duck, 1990) |
| Brain (white matter) | 3600 J/kg/K | (Collins et al., 2004) |
| Brain | 3630 J/kg/K | (Baumgartner et al., 2025):<br>(McIntosh & Anderson, 2010; Williams & Leggett, 1989) |
| Brain (gray matter) | 3680 J/kg/K | (Collins et al., 2004) |
| Brain (gray matter; human) | 3680 J/kg/K | (Duck, 1990) |
| Brain (gray matter) | 3695.8 J/kg/K | (Baumgartner et al., 2025):<br>(Bernardi et al., 2003; Duck, 1990; McIntosh & Anderson, 2010) |
| Brain (gray matter; human) | 3718 J/kg/K | (McIntosh & Anderson, 2010) |

| PERFUSION (mL/min/kg) | | |
|---|---|---|
| **Tissue** | **Value** | **Reference** |
| Brain (white matter; human) | 196 mL/min/kg | (McIntosh & Anderson, 2010) |
| Brain (white matter) | 200 mL/min/kg | (Biagi et al., 2007) |
| Brain (white matter) | 230–250 mL/min/kg | (Parkes et al., 2004) |
| Brain (gray matter) | 231–287 mL/min/kg | (Liu et al., 2012) |
| Brain (white matter) | 237 mL/min/kg | (Collins et al., 2004) [15] |
| Brain (white matter) | 320-440 mL/min/kg | (Muer et al., 2024) |
| Brain | 450-620 mL/min/kg | (Muer et al., 2024) |
| Brain (gray matter) | 474 mL/min/kg | (Petersen et al., 2010) |
| Brain (human) | 520 mL/min/kg | (McIntosh & Anderson, 2010) |
| Brain | 530-620 mL/min/kg | (Parkes et al., 2004) |
| Brain (gray matter) | 540-750 mL/min/kg | (Muer et al., 2024) |
| Brain | 560 mL/min/kg | (Williams & Leggett, 1989) |
| Brain (gray matter) | 580-680 mL/min/kg | (Parkes et al., 2004) |
| Brain (gray matter) | 580 mL/min/kg | (Biagi et al., 2007) |
| Brain (gray matter) | 610 mL/min/kg | (Zhang et al., 2018) |
| Brain (gray matter) | 671 mL/min/kg | (Collins et al., 2004) |
| Brain (gray matter; human) | 809 mL/min/kg | (McIntosh & Anderson, 2010) |

[15] Collins et al. (2004) compiled values from the literature and give no specific source for each value.

### Supplementary Table S1c. Water

| SOUND SPEED (m/s) | |
|---|---|
| **Value** | **Reference** |
| 1482.3 m/s | (Baumgartner et al., 2025): (Kyriakou, 2015) |

| DENSITY (kg/m³) | |
|---|---|
| **Value** | **Reference** |
| 994.04 kg/m³ | (Baumgartner et al., 2025) |

| ATTENUATION (dB/cm) | | | |
|---|---|---|---|
| **dB/cm @ 1 MHz** | **Exponent b** | **dB/cm @ 0.5 MHz** | **Reference** |
| 0.00191 dB/cm/MHz | 2 | 0.0004775 dB/cm | (Kaye & Laby, 1995): physical minimum |
| 0.0022 dB/cm/MHz | 2 | 0.00055 dB/cm | (Baumgartner et al., 2025): (Kyriakou, 2015) |
| 0.002 dB/cm/MHz | 2 | 0.0005 dB/cm | (Duck, 1990) |

| THERMAL CONDUCTIVITY (W/m/K) | |
|---|---|
| **Value** | **Reference** |
| 0.6045 W/m/K | (Baumgartner et al., 2025) |

| SPECIFIC HEAT (J/kg/K) | |
|---|---|
| **Value** | **Reference** |
| 4178 J/kg/K | (Baumgartner et al., 2025) |
| 4190 J/kg/K | (Giering et al., 1995) |

| PERFUSION (mL/min/kg) | |
|---|---|
| **Value** | **Reference** |
| 0 mL/min/kg | (Baumgartner et al., 2025) |

| ABSORPTION (dB/cm @ 500 kHz) | |
|---|---|
| **Value** | **Reference** |
| 0.00055 dB/cm (100%) | (Hill et al., 2004) |

**Supplementary Table S2.** Brain and skull properties of the literature setups and the emulated toolbox defaults, as simulated in the toolbox comparison (Figure 5). Attenuation α and absorption β are given at 500 kHz; α sets acoustic propagation and β sets heat deposition. Temp-H / Temp-L: per property, the LOW or HIGH bound of Table 1 that raised / lowered temperature rise in both skull and brain in the one-at-a-time analysis (skull sound speed: skull only). Trans-H / Trans-L: per acoustic property, the bound that maximizes / minimizes transcranial transmission; absorption and thermal properties are not specified as these setups bracket intensity only. * Emulated in PRESTUS from the tool defaults. k-Plan skull sound speed follows its density mapping (1.33 ρ + 166.7) at the thermal-simulation density of 1850 kg/m³. Water (all setups): c = 1482 m/s, ρ = 994 kg/m³, α = 0.00055 dB/cm at 500 kHz (y = 2), β = 0.00055 dB/cm, κ = 0.6 W/m/K, $c_p$ = 4178 J/kg/K, no perfusion.

| | **Property** | **Unit** | **Temp-H** | **Temp-L** | **Trans-H** | **Trans-L** | **PRESTUS** v0.6.1 | **k-Plan*** v1.2.1 | **BabelBrain*** v0.8.8 |
|---|---|---|---|---|---|---|---|---|---|
| Brain | Sound speed c | m/s | 1573 | 1520 | 1520 | 1573 | 1546 | 1550 | 1562 |
| | Density ρ | kg/m³ | 1040 | 1050 | 1050 | 1040 | 1046 | 1045 | 1041 |
| | Attenuation α at 500 kHz | dB/cm | 0.492 | 0.154 | 0.154 | 0.492 | 0.257 | 0.26 | 0.24 |
| | Absorption β at 500 kHz | dB/cm | 0.492 | 0.0447 | - | - | 0.257 | 0.26 | 0.204 |
| | Absorbed fraction β/α | - | 1.00 | 0.29 | - | - | 1.00 | 1.00 | 0.85 |
| | Frequency power law y | - | 1 | 1 | 1 | 1 | 1.2 | 1 | 1 |
| | Thermal conductivity κ | W/m/K | 0.48 | 0.55 | - | - | 0.51 | 0.51 | 0.51 |
| | Specific heat capacity $c_p$ | J/kg/K | 3525 | 3718 | - | - | 3630 | 3600 | 3630 |
| | Perfusion ω | mL/min/kg | 196 | 809 | - | - | 559 | 0 | 559 |
| Skull | Sound speed c | m/s | 2871 | 2190 | 2190 | 2871 | 2800 | 2627 | 2800 |
| | Density ρ | kg/m³ | 1080 | 1991 | 1080 | 1991 | 1850 | 1850 | 1850 |
| | Attenuation α at 500 kHz | dB/cm | 4.02 | 11.7 | 4.02 | 11.7 | 6.65 | 6.65 | 6.65 |
| | Absorption β at 500 kHz | dB/cm | 4.02 | 1.05 | - | - | 1.86 | 6.65 | 1.08 |
| | Absorbed fraction β/α | - | 1.00 | 0.09 | - | - | 0.28 | 1.00 | 0.16 |
| | Frequency power law y | - | 1 | 1 | 1 | 1 | 1 | 1 | 1 |
| | Thermal conductivity κ | W/m/K | 0.3 | 0.36 | - | - | 0.32 | 0.32 | 0.315 |
| | Specific heat capacity $c_p$ | J/kg/K | 826 | 2524 | - | - | 1793 | 1300 | 1793 |
| | Perfusion ω | mL/min/kg | 0 | 30 | - | - | 20 | 0 | 20 |

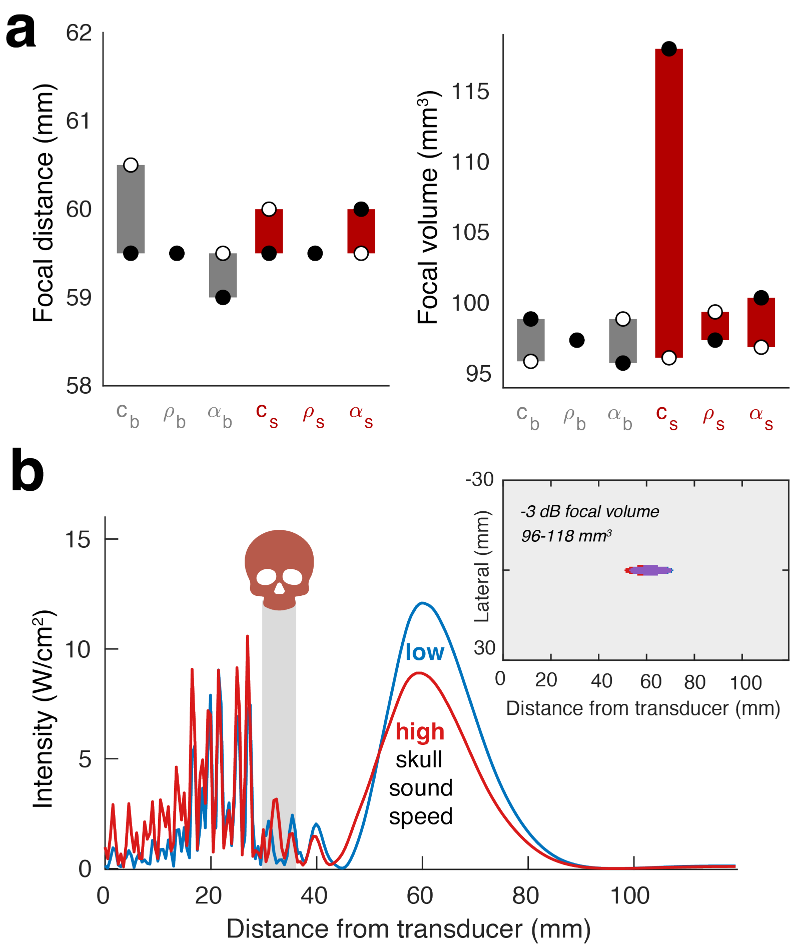


**Supplementary Figure 1. Focal position and volume are largely robust to acoustic property assumptions.** (a) Focal distance from the transducer (left) and −3 dB focal volume (right) in the skull benchmark when one acoustic property of brain (subscript b, gray) or skull (subscript s, red) is set to its LOW (white dot) or HIGH (black dot) bound, with all other properties at reference values; bars span the two outcomes. c, sound speed; ρ, density; α, attenuation coefficient. No property shifts the focus by more than 1 mm. Skull sound speed dominates the focal volume (96–118 mm³); the other properties change it by about 4 mm³ or less. (b) On-axis intensity for LOW (blue, 2190 m/s) and HIGH (red, 2871 m/s) skull sound speed. The gray band marks the skull. Inset: −3 dB focal volumes of both settings (blue, LOW only; red, HIGH only; purple, overlap).

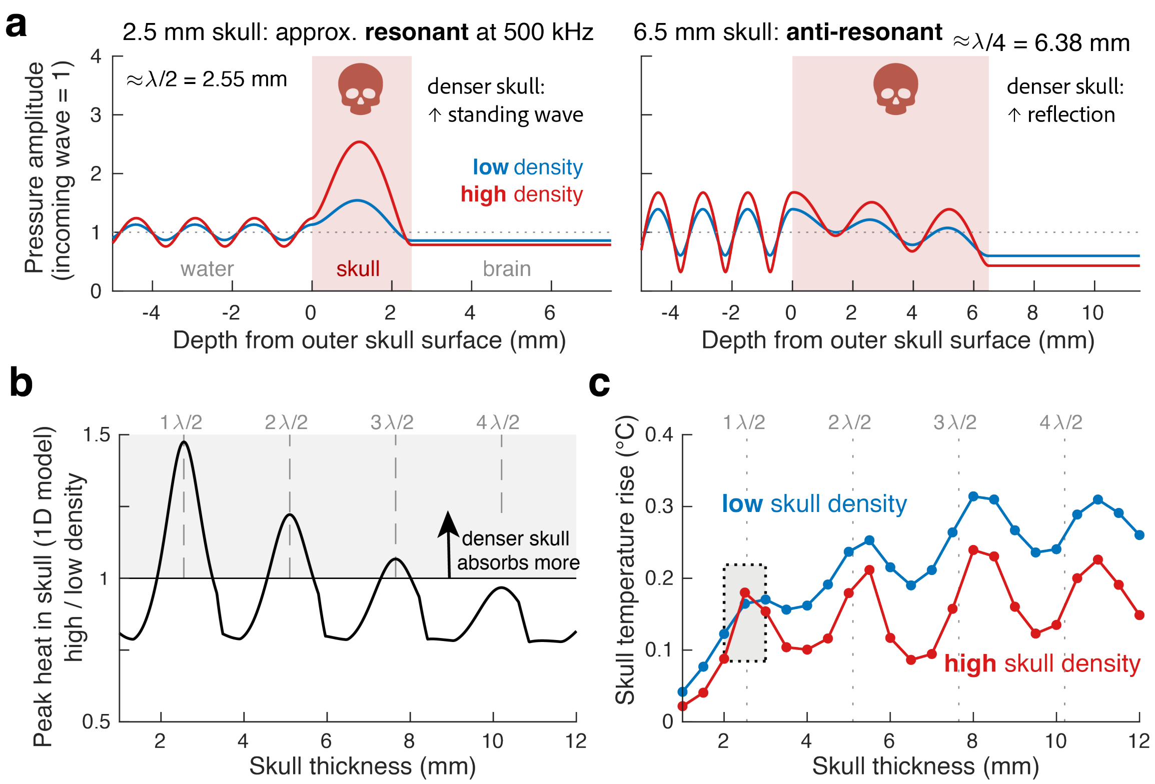


**Supplementary Figure 2. Skull density has opposite effects on skull heating at resonant and non-resonant skull thicknesses.** (a) Schematic 1D plane-wave model (water | skull plate | brain; 500 kHz, normal incidence) for LOW and HIGH skull density (1080 and 1991 kg/m³), with skull sound speed (2550 m/s), attenuation and absorption at reference values. The analytic steady-state pressure amplitude is shown relative to the incident wave; shading marks the skull. Left: the 2.5 mm skull is close to resonant (λ/2 = 2.55 mm, with λ the wavelength in bone). Repeated reflections inside the plate build a standing wave that grows with the impedance contrast between bone and water, so the denser skull holds a stronger field and absorbs more heat. Right: the 6.5 mm skull is close to anti-resonant (5λ/4 = 6.38 mm); the denser skull reflects more of the wave and absorbs less. (b) Model prediction: HIGH/LOW density ratio of peak heat deposition in the skull as a function of skull thickness. The denser skull absorbs more near half-wave resonances (dashed lines at multiples of λ/2) and less elsewhere. (c) Thermal simulations: skull temperature rise against skull thickness in PRESTUS (2D axisymmetric, 20 s sonication, all other properties at reference values). The gray bar marks the 2.5 mm benchmark skull. The high-density curve approaches the low-density curve near the resonances but exceeds it only at the first. With specific heat capacity held constant, density also scales the volumetric heat capacity of bone, so a denser skull warms less for the same deposited heat, and only the largest resonance gain outweighs this.

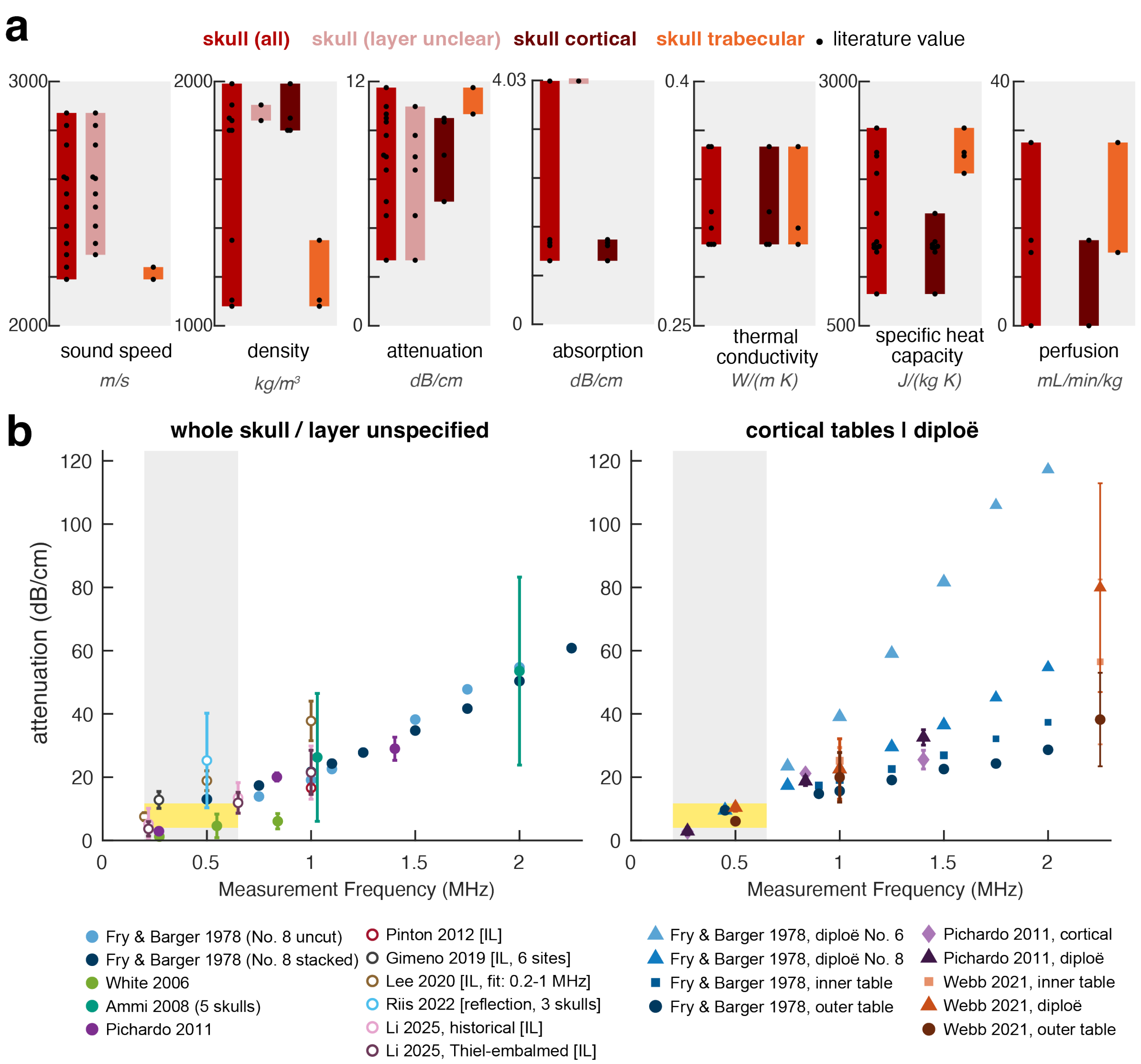


**Supplementary Figure 3. Skull property estimates by layer.** (a) Literature values of skull acoustic and thermal properties (Supplementary Table S1), pooled and split by layer (layer unclear, cortical, trabecular). Bars span the range of values; dots mark individual values; n/a, no values for that layer. Attenuation and absorption coefficient in dB/cm at 0.5 MHz. Cortical and trabecular values do not overlap for density, attenuation and specific heat capacity. A dynamic visualization is provided in the ITRUSST Tissue Properties database (itrusst.github.io/itrusst_properties/). (b) Skull attenuation as a function of measurement frequency for whole/unspecified skull (left) and for cortical table and diploë measurements (right). Points are the values measured or fitted in each study at each frequency. Open markers indicate values that include reflection losses: insertion losses divided by the reported skull thickness ([IL]) and Riis 2022, whose reflection term is a fitted intercept. For studies reporting one value per skull or site (Ammi et al., 2008; Gimeno et al., 2019; Riis et al., 2022), points show the mean ± SD across skulls or sites. Gray shading marks frequencies common in transcranial ultrasound stimulation (0.2–0.65 MHz); the dotted line marks the simulation frequency (0.5 MHz). Yellow shading marks the Table 1 skull attenuation range (LOW–HIGH) at 0.5 MHz, and the dashed black line the reference. Reported values are from (Ammi et al., 2008; Fry & Barger, 1978; Gimeno et al., 2019; Lee et al., 2020; Li et al., 2025; Pichardo et al., 2011; Pinton et al., 2012; Riis et al., 2022; Webb et al., 2021; P. J. White et al., 2006).
